# Kinetics of RNA and RNA:DNA hybrid strand displacement

**DOI:** 10.1101/2021.04.01.438146

**Authors:** Hao Liu, Fan Hong, Francesca Smith, John Goertz, Thomas E. Ouldridge, Hao Yan, Petr Šulc

## Abstract

In dynamic nucleic acids nanotechnology, strand displacement is a widely used mechanism where one strand from a hybridized duplex is exchanged with an invading strand which binds to a toehold, a single-stranded region on the duplex. With proper design and kinetic control, strand displacement is used to perform logic operations on molecular level to trigger the conformational change in nanostructures, initiate cascaded reactions, or even for in vivo diagnostics and treatments. While systematic experimental studies have been carried out to probe the kinetics of strand displacement in DNA, there has not been a comparable systematic work done for RNA or RNA-DNA hybrid systems. Here, we experimentally study how toehold length, toehold location (5′ or 3′ end of the strand) and mismatches influence the strand displacement kinetics. Through comparing the reaction rates, combined with previous theoretical studies, we observed reaction acceleration with increasing toehold length and placement of toehold at 5′end of the substrate. We find that mismatches closer to the interface of toehold and duplex slow down the reaction more than remote mismatches. Comparison of RNA displacement and DNA displacement with hybrid displacement (RNA invading DNA or DNA invading RNA) is in part explainable by the thermodynamic stabilities of the respective toehold regions, but also suggest that the rearrangement from B-form to A-form helix in case of RNA invading DNA might play a role in the kinetics. The measured kinetics of toehold-mediated strand displacement will be important in understanding and construction of more complex dynamic nucleic acid systems.

## I. INTRODUCTION

DNA and RNA play essential roles in all living systems. Beside their importance in biological systems, they have also become essential in nanotechnology applications due to their structural diversity and programmability enabled by base pairing between complementary strands. In the past few decades, the fields of DNA and RNA nanotechnology led to design of increasingly complex nanostructures and devices self-assembled out of DNA and/or RNA, with applications ranging from photonics and nanofabrication to drug delivery, medical treatment and diagnostics [1–4].

A key mechanism in the active nucleic acid nanotechnology is the strand displacement reaction [5, 6]. Starting from the pioneering work of DNA nanotweezers done by Yurke et al [7], toehold mediated strand displacement has been employed to construct complicated nucleic acid system. The process has found a wide range of applications that include diagnostics [3], reconfigurable and dynamic nanostructures [8, 9], and nanomotors [10] to molecular computation [11–13]. It is also likely involved in interactions between RNA molecules in biological systems [14]. Recent experimental studies have also shown that internal strand displacement (where one part of RNA strand displaces other regions of the same RNA strand) is also likely involved in SRP RNA kinetic folding pathway [15].

The strand displacement reaction involves an invading strand (invader), which binds to a single-stranded region (toehold) on a complementary strand (substrate), displacing a partially complementary strand (incumbent). Readout is often accomplished via an additional duplex (reporter) with fluorophore-modified ends that are separated by a second strand displacement reaction between it and the incumbent, as illustrated in Fig. 1. In short, the strand displacement reaction describes the process in which the invader binds to the complementary substrate strand previously bound to the incumbent strand and hence the latter is released. This hybridization reaction exchanges the incumbent strand from the duplex with the new invader and could be therefore used to initiate the conformational change or cascaded reactions, and it was shown that combination of these reactions can realize complex logic functions, where the input is presence or absence of a particular strand [13]. The presence of toehold increases the probability to capture the invader and hence speeds up the kinetics of the removal of the incumbent strand by the invader, with the speed increasing with longer toeholds [6]. Toehold mediated strand displacement (TMSD) reaction has become a popular choice for researchers to construct dynamic structures.

**FIG. 1.**
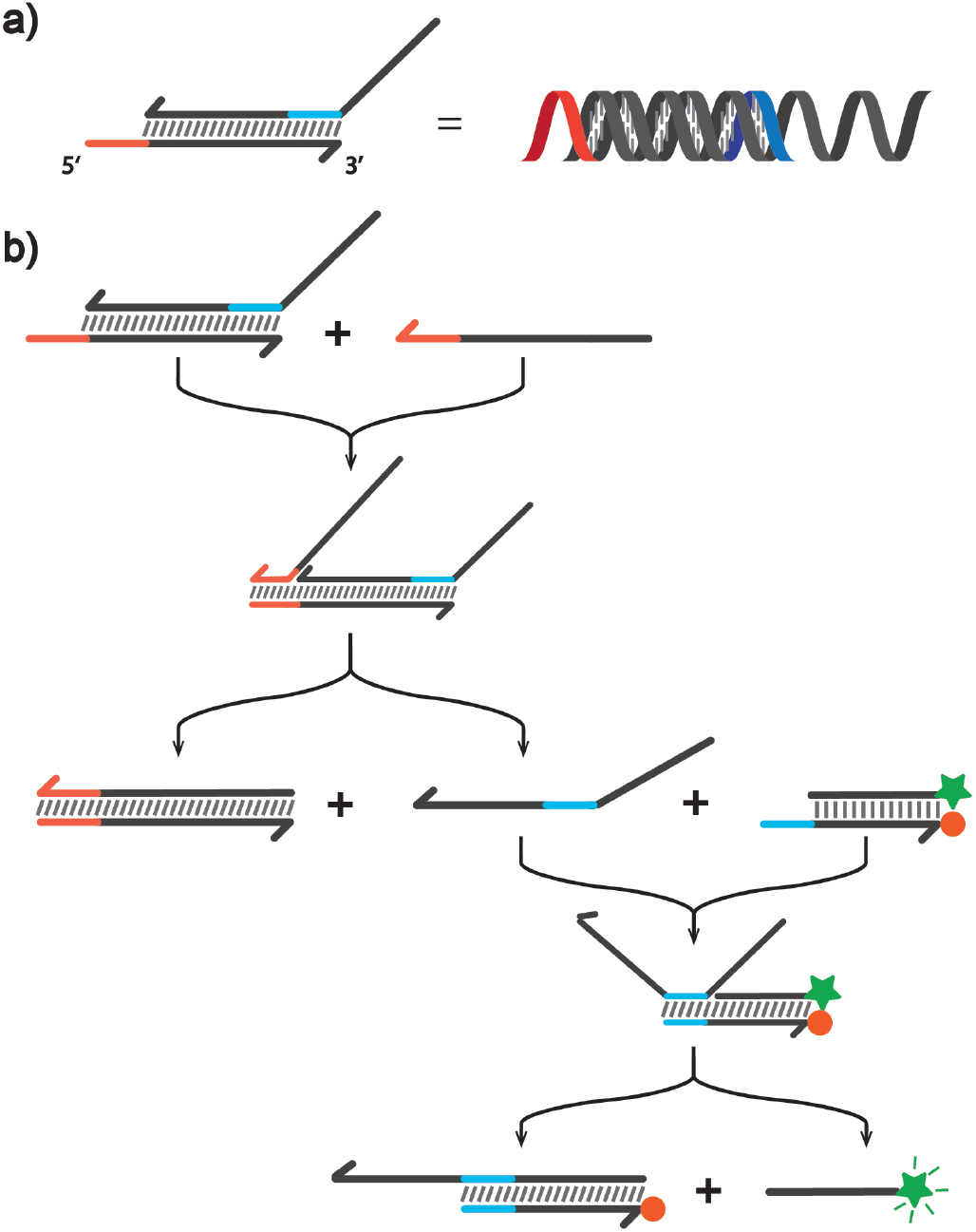
Schematic illustration of the experimental design of RNA strand displacement. a) RNA strand is represented using half headed arrow with 3 end being the arrow side.RNA Duplex is represented with gray slash between two RNA strand and the toehold is colored with orange and blue. b) Displacement of the RNA incumbent strand triggers the secondary strand displacement of FAM labeled fluorescent strand which was quenched by the adjacent TAMRA labelled strand through FRET. Fluorescent signal could be therefore observed within the FAM emission wavelength.

In spite of the extensive usage of DNA in nanodevices that use strand displacement, RNA and RNA:DNA hybrid stand displacement is also of great interest due to its potential in *in vivo* application, ranging from synthetic biology to diagnostics [16]. Moreover, the RNA:DNA hybrid strand displacement is also speculated to be be involved in the gene editing process of CRISPR-Cas9 system, where the formation of R-loop requires the base flipping in target DNA double strand for crRNA to bind with the assistance of nearby amino acids[17, 18]. The transition of DNA-DNA duplex to RNA-DNA hybrid duplex could be rationalized with strand displacement process [19, 20]. Further experimental proofs are necessary to verify the hypothesis while more insights could also be gained through studying the RNA:DNA hybrid strand displacement system.

The efforts to understand the mechanism of strand displacement could be dated to the initial studies of branch migration [21, 22]. Yurke and Mills [23] first identified the phenomena of exponential acceleration of the rate constant with the increasing toehold length with the study of kinetics. Zhang and Winfree [6] then studied the mechanism of the DNA strand displacement through systematic kinetics experiment which revealed that saturated toehold length for reaction acceleration was 6 and proposed a quantitative model of DNA strand displacement with varying toehold length. Srinivas et al [24] proposed an approximate empirical model called “Intuitive Energy Landscape” (IEL) model through the computational studies which included the branch migration with more physical details. A coarse-grained model of DNA [25] was also used to study strand displacement reactions through simulation which provides more structural and physical understanding [24]. Furthermore, the effects of mismatches between the invader and the substrate on the displacement kinetics of DNA TMSD has been studied in Ref. [26, 27]. In the case of “mismatch repair” the mismatches were placed between the incumbent and substrate and therefore corrected by the displacement [27]. In the case of “mismatch introduction” the mismatches are between the invader and substrate and the displacement generates mismatched sites. In both cases the kinetics are highly dependent on the position of the mismatch [26–28].

Systematic study of RNA TMSD was performed *in silico* with oxRNA [29], a coarse-grained model of RNA. Qualitatively similar dynamic behavior was observed compared to DNA strand displacement when the toehold length was varied [30], with exponential speed-up of TMSD rate with increasing toehold length until the speed is saturated. Additionally, toehold location (5′ or 3′) was also predicted to affect the reaction rate due to an extra cross-stacking interaction between the invader and substrate strands when toehold is located at the 5′ end, which thus speeds up the reaction. Recently, an experimental study has been carried out where an RNA invades DNA duplex [31]. They compared DNA and RNA single strand invading DNA duplex with long toehold, and found that location of toehold as well as the toehold sequence affect the kinetics of the displacement.

Here we systematically study, for the first time in experiment, the RNA strand displacement, as well as hybrid systems with DNA invading RNA duplex and RNA invading DNA duplex. We further examine the effects of mismatches between the RNA invader and the substrate, as well as the kinetics of toehold positioned either at 5′or 3′ end of a substrate.

The quantitative analysis of the kinetics result along with the proper modelling will be beneficial for further designs of RNA-involved strand displacement systems especially in the fields of synthetic biology and RNA nanotechnology.

## II. MATERIALS AND METHODS

### A. Sequence Design

The sequence for the strand displacement characterization is designed using NUPACK [32] to minimize undesired secondary structures. For the experiments where we compare the 3′ and 5′ toeholds, we designed the toehold sequences to have the same free-energy of binding to the invader and the first and last bases of the branch migration are set to be Gs to minimize the change of toehold stacking energy. For example, the 5 toehold is 5GACCAG–, the corresponding 3 toehold is –GACCAG-3, and the stacking base for the 5 toehold and 3 toehold are both Gs. All the sequences of RNA and DNA strands that were used in the experiments are listed in Table S1 - S4 and the NUPACK prediction on each strand is shown in Figure S1 - S3.

### B. Experiment Design

A schematic illustration is shown in Figure 2. To study the mismatch effects on strand displacement, six typical positions of the sequence, distributed in both branch migration range and toehold range, are selected and changed to another nucleotide that will not pair to the corresponding nucleotide on the other strand. For the study of both mismatches and toehold length effects, the toehold is located at the 5 end of the substrate. The toehold length is varied from 1 nt to 6 nt. Additionally, experiments with 3 end toehold are conducted for toehold lengths 1, 3, and 6 nucleotides. For RNA-DNA hybrid systems, we perform experiments where a) an RNA invader displaces a DNA incumbent strand from a DNA substrate, and b) a DNA invader displaces an RNA incumbent strand from an RNA duplex. As a control, the strand displacement for pure DNA system were designed to study the kinetics in which toehold are also varied with typical lengths for comparison.

**FIG. 2.**
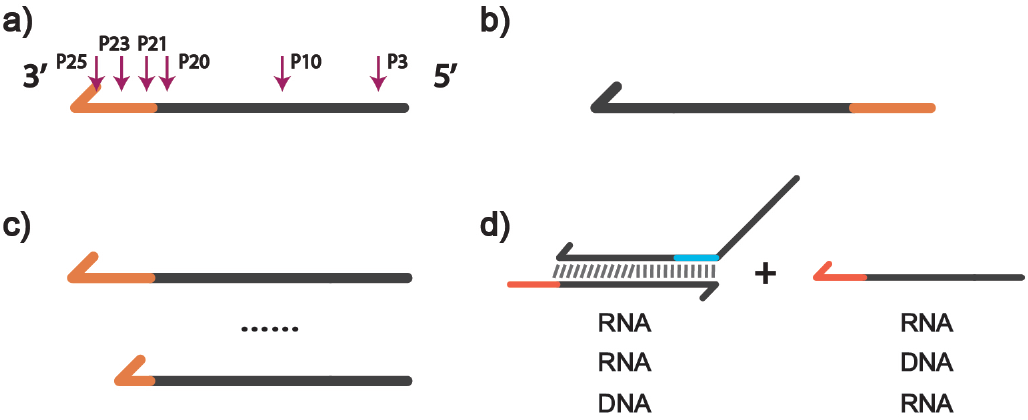
The design of the invader strand in different experiments. a) Mutation is located at a position 3, 10, 20, 21, 23, or 25. The first three positions are located in the branch migration region and the last three placed within the toehold. We only consider experiments where a single mutation is present. b) Toehold located at the 3 end of the substrate strand and correspondingly at the 5 end of the invader. c) Toehold length was varied from 1 nucleotide (nt) to 6 nt on the invader. d) Combinations for RNA-DNA hybrid experiments, where additional to RNA strands invading RNA duplex, we consider RNA invading DNA duplex, DNA invading RNA duplex, and DNA invading DNA duplex, where we keep the sequence and toehold length identical across all experiments

Although our experiments study RNA strand displacement, the reporter strand is DNA-based due to the fact that RNA strand displacement and signal readout are two separate processes and the latter will be fitted individually for rate constants, so the stability provided by DNA will be preferred.

### C. Substrate Preparation

Nucleic acid oligonucleotides used in here were purchased from Integrated DNA Technologies (IDT), and purified through denaturing poly-acrylamide gel electrophoresis. Where applicable, fluorophores were attached by IDT as well. The measured absorbance at 260 nm using Thermo Scientific NanoDrop 2000 along with the calculated extinction coefficient were used to determine concentrations. All double-stranded gates were prepared through annealing processes in a thermocycler (Eppendorf) with different programs in 1 X TAE Mg^2+^ buffer (Tris base 40 mM, acetic acid 20 mM, ED-TANa212H2O 2 mM, (CH3COO)2Mg4H2O 12.5 mM).

The RNA samples were annealed at 65°C for 5 minutes and then brought down from 20°C with a constant rate of 1°C/min while for DNA samples, after 5 minutes of annealing, they were programmed to be cooled down to 20°C with the same rate.

### D. Fluorimetry Experiments

The fluorescent kinetics over the strand displacement time was monitored with a Nanolog fluorometer (Horiba Jobin Yvon). All kinetic experiments were performed at 25°C in 1 X TAE Mg^2+^ buffer in a lid-covered Hellma Analytics cuvette. To minimize the non-specific cohesion of nucleic acids under low concentration to the tubes wall, the sample to be measured with final concentration (60 nM) was diluted in the cuvette from 2 *μ*M stock solution. To initiate the reaction, incumbent was added to the reporter solution for the single gate one and incumbent-bottom gate was added to the mixture of reporter and invader for the double gate reaction.

Before the initiation of reaction, signals and sample’s temperature were allowed to stabilize for at least 300s. The addition of strands was kept with the same fashion by rapidly pipetting the solution and data points collected at the first 12s were discarded due to the influence of pipetting. Cuvettes were thoroughly cleaned subsequently by Milli-Q purified water and pure ethanol and allowed to be fully dried for next measurement.

The parameters settings of the fluorescence measurements were as follows: 495 nm excitation, 1 nm excitation slit, 520 nm emission, 10 nm emission slit. Specifically, a 1 nm excitation slit was chosen to reduce photobleaching of the dye molecules while a 10 nm emission slit guaranteed a proper signal level to be observed. The signal was collected from 0 to 5000 s with 0.5 s integration time and 3 s intervals. Kinetics measurements were repeated 3 times.

### E. Data processing

The fluorescent signal was normalized with respect to the positive control (60nM fluorescently-labelled strand) and the initial fluorescent signal at time (t = 0), resulting in a time-dependent fluorescent signal between 0 and 1.

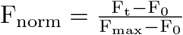

Above equation describes the normalization procedure, in which F_norm_ represents the normalized fluorescence; F_t_ the raw fluorescence data; F_0_ the initial fluorescence; and F_max_ the maximum achievable fluorescence, as indicated by the positive control. The positive control is done by observing the fluorescence change of 60 nM reporter strand only for 5000 seconds. This gave a normalized fluorescent signal value between 0 and 1. Specifically, the first point does not start from 0 second as we removed the incorrectly changed signal part due to the pipetting and as a result, subtracting the data-set with the number corresponding to the initial point introduces extra error and lowers down the final signal level it is able to reach. However, we do not observe significant difference on the resulted rate constant through introducing such normalization method and for fitting’s convenience we keep it for the consistency. Based on these normalized fluorescent signals we observed that most of the experimental systems reached equilibrium with fluorescent signal smaller than the maximum fluorescence. Following the approach previously employed for DNA [6], we introduced an additional parameter *α*, which represents a scaling factor capturing the proportion of the maximum fluorescence signal achieved at equilibrium. Thus, intuitively, *α* represents the proportion of the *c*_0_ = 60 nM initial reactant concentration which reacted.

### F. Data fitting

While different versions of fitting of TMSD reaction is studied in this paper (see Supp. Mat.), a general case can be represented as below:

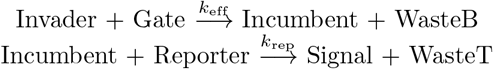

The reaction kinetics is assumed to be of second order and a notation for the related reaction species is used as: Invader (V), Gate (G), Incumbent (I), Reporter (R) and Signal (S). *k*_eff_ and *k*_rep_ are therefore the rate constants for the displacements of the incumbent by invader and the signal by incumbent respectively. The differential relation between each reaction species can be modelled as follows:

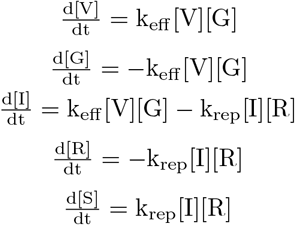

with initial conditions [I](0) = [S](0) = [R](0) = c_0_*α* and others set to 0. The best-fit *k*_rep_ and *α* parameters for the reporter characterization experiments are determined (see Supplementary Material) and subsequently with the known *k*_rep_ using the outlined ODEs we fit values for *k*_eff_ and *α* to match the measured fluorescent signal. We constrain the upper and lower bounds *k*_rep_ and *k*_eff_ to 1 to 7 and 1 to 8, respectively.

The fitting method employed here corresponds to the method adapted from [6]. We tried the method from [26] (see Supplementary Table S6) for fitting as well. However, the method from [26] assumes that the reporter reaction is much faster than the original strand displacement reaction, which is not fully justified for our system, hence we report here the fits from sets of equations described above.

### G. RNA strand handling and secondary structure

Compared to the study on DNA strand displacement[6, 26, 28, 33], the inherent instability of RNA due to hydroxyl group as well as possibility of degradation from ubiquitous RNases can introduce experimental artifacts. Furthermore, secondary structure formed in RNA is more stable than it would be for the corresponding DNA sequence due to larger stability of RNA canonical base pairs. Extra attention has been paid to both the sequence design and experimental operation to avoid factors that will affect the result and conclusion. However, as indicated by native gel electrophoresis (Figure S4), we found for certain sequences that RNA secondary structure still forms in our experiment even if it is not predicted by NUPACK (Figure S1). It was confirmed from both kinetic curves measured (Figure S5) and fitted rate constants (Table S6) that the formation of secondary structure slows down the reaction significantly, and we discarded toehold sequences that form secondary structure from our analysis in the main text. To prevent degradation, RNA sample stock were properly stored in 80°C and handled with RNase free water and pipette tips.

Control experiments (Figure S6) were conducted with RNase inhibitor added to confirm during the fluorimetry measurement RNA is not degraded. Non-denaturing gel electrophoresis (Figure S4) serves as a method to identify whether RNA is degraded or not by checking the band condition and single band without the typical smear proves that within the sensitivity of gel electrophoresis the RNA samples used have not degraded significantly. While it does not rule out the potential degradation concerns, the clear trend shown from the curves and the limited errorbar of them proves that the efforts made to avoid unwanted factors are effective. However, the fitting method adopted does not work for all the measurements as not all reactions reach to the equilibrium within the measuring time, and also with slower reaction rate, the second order reaction assumption might not apply due to both the slightly different biophysical process and relatively more significant noise. Specifically, the experiments are done with all the reacting species have the same concentration, and the strand displacement reaction therefore saturate in a relative slow way, resulting the fact that not all the curves shown reach the expected equilibrium. The intrinsic slow reactions are in this case more susceptible which reduces the precision of the developed fitting model on producing a rate constant. The over-estimate for slow reactions with the fitting method adopted in this research poses problems on evaluating the rate constants in an absolute scale.

## III. RESULTS AND DISCUSSION

We design a set of experiments to measure the TMSD reaction kinetics. As schematically outlined in Fig. 1, the displaced strand then binds to a DNA reporter complex that produces a fluorescent signal. To study the kinetics of RNA strand displacement, we consider systems of variable toehold lengths (from 1 to 6 nt) placed either at 3′ or 5′ end of the substrate. We additionally test systems where the invading RNA strand has a single mismatch with the substrate strand. We consider six different positions of the mismatch location. We further study the cases of hybrid DNA:RNA strand displacement, where either the DNA invader displaces incumbent RNA strand bound to an RNA substrate, or an RNA invader displaces DNA incumbent from a DNA substrate. The systems studied in experiment are schematically shown in Fig. 2. The details of the experimental setup and the data fitting procedure is described in Methods section and in the Supplementary Material. The experimental results for the respective systems studied are provided below and the fitted rates of TMSD reactions for all the systems studied are shown in Table I.

**TABLE I.**
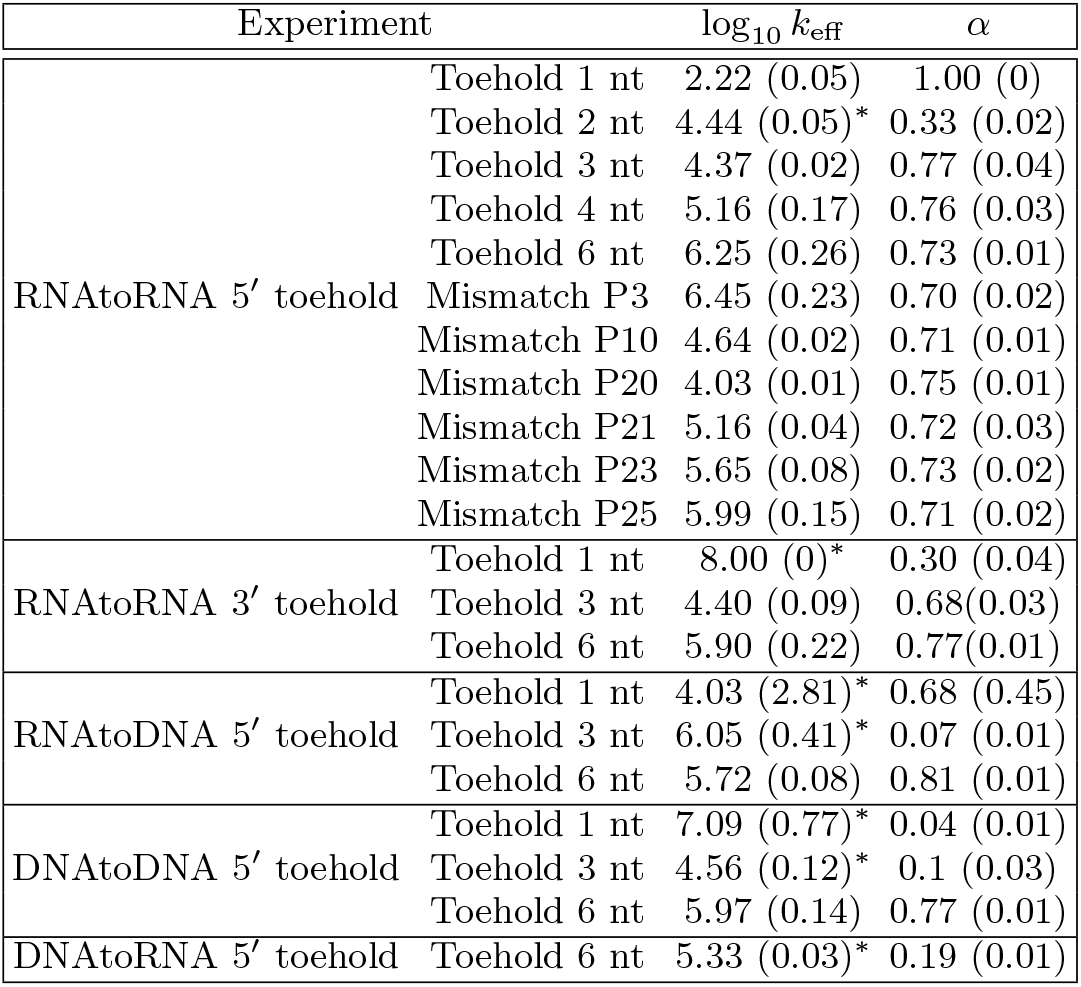
Fitted rate constants to the experimental data. Rate constants with low *α* value are considered to be incorrectly fitted and they are labelled with asterisk symbol. All the rate constants and the associated alpha are shown as the average of the fitting results of the three replicas and the calculated standard deviation is shown in the bracket.

| Experiment | | $\log_{10} k_{\text{eff}}$ | $\alpha$ |
| --- | --- | --- | --- |
| RNAtoRNA 5' toehold | Toehold 1 nt | 2.22 (0.05) | 1.00 (0) |
|  | Toehold 2 nt | 4.44 (0.05)* | 0.33 (0.02) |
|  | Toehold 3 nt | 4.37 (0.02) | 0.77 (0.04) |
|  | Toehold 4 nt | 5.16 (0.17) | 0.76 (0.03) |
|  | Toehold 6 nt | 6.25 (0.26) | 0.73 (0.01) |
|  | Mismatch P3 | 6.45 (0.23) | 0.70 (0.02) |
|  | Mismatch P10 | 4.64 (0.02) | 0.71 (0.01) |
|  | Mismatch P20 | 4.03 (0.01) | 0.75 (0.01) |
|  | Mismatch P21 | 5.16 (0.04) | 0.72 (0.03) |
|  | Mismatch P23 | 5.65 (0.08) | 0.73 (0.02) |
|  | Mismatch P25 | 5.99 (0.15) | 0.71 (0.02) |
| RNAtoRNA 3' toehold | Toehold 1 nt | 8.00 (0)* | 0.30 (0.04) |
|  | Toehold 3 nt | 4.40 (0.09) | 0.68(0.03) |
|  | Toehold 6 nt | 5.90 (0.22) | 0.77(0.01) |
| RNAtoDNA 5' toehold | Toehold 1 nt | 4.03 (2.81)* | 0.68 (0.45) |
|  | Toehold 3 nt | 6.05 (0.41)* | 0.07 (0.01) |
|  | Toehold 6 nt | 5.72 (0.08) | 0.81 (0.01) |
| DNAtoDNA 5' toehold | Toehold 1 nt | 7.09 (0.77)* | 0.04 (0.01) |
|  | Toehold 3 nt | 4.56 (0.12)* | 0.1 (0.03) |
|  | Toehold 6 nt | 5.97 (0.14) | 0.77 (0.01) |
| DNAtoRNA 5' toehold | Toehold 6 nt | 5.33 (0.03)* | 0.19 (0.01) |

### A. Effects of toehold length and location

We first studied the effects of the toehold length on the TMSD kinetics with the toehold placed at the 5′ end of the substrate. We considered toehold lengths 1, 2, 3, 4 and 6 nt. As shown in the Fig. 3a and Table I, a clear trend emerges that increasing the toehold length accelerates the reaction rate for both the 5’ end toehold location as well as 3’ end toehold location. We observe that for the toeholds longer than 2 nts, the fitted reaction rate increases by about a factor of 6 to 10 with increasing the toehold length by one base pair (Table I).

**FIG. 3.**
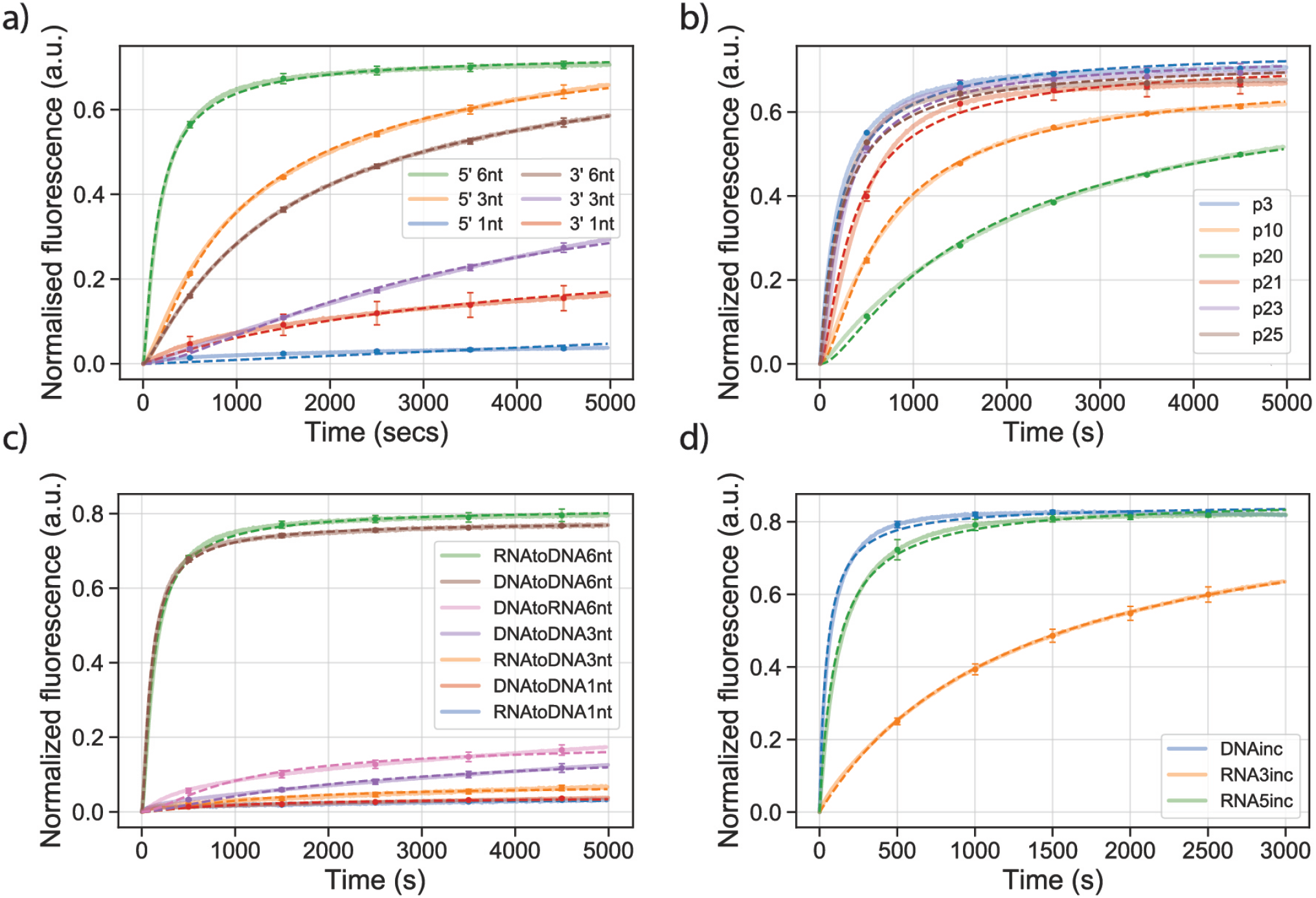
Summary of the kinetic profiles of RNA strand displacement experiment. a) Toehold located at either 5 or 3’ end with length 1, 3, 6 nt. b) Mismatches placed in different positions selected from both branch migration range and toehold range. For example, P3 in the legend means the third base pair counted from 5’ forms mismatch. RNA:RNA, DNA:DNA and RNA:DNA hybrid strand displacement curves with toehold length being 1, 3 and 6nt. In the legend, “RNAtoDNA” means RNA invader displaces DNA gate and similarly the meaning for others can be derived. d) The reaction involves the incumbent strand and the reporter to characterize *k*_*rep*_. In the legend, “RNA5inc” means it is a RNA incumbent strand with the toehold placed at the 5 end. The fluorescent signals are normalized according to the protocol described in data processing section. All curves, unless specified, correspond to reactions with toehold placed at 5’ end and length being 6 nt. Each curve that represents the trajectory of the fluorescent kinetics is shown as thick and transparent while the dotted line in the same color represents the fitting result given by the fitted model. The error bars represent the standard deviation calculated from 3 replicate measurements and they are shown at 500s, 1500s, 2500s, 3500s and 4500s respectively. They are not shown if the errorbar is smaller than the symbol itself.

This behavior is in agreement with previous experimental measurements for DNA [6, 24] as well as simulation studies of RNA TMSD [30]. For shorter toeholds, the probability of the invader binding to the substrate decreases as the weaker interaction strength leads to frequent binding and unbinding of the invader from the toehold. Once the free energy of binding to the toehold becomes sufficiently strong, the rate of displacement saturates, as the probability of binding is so small that the invader always completes the strand displacement after binding to the toehold.

We note that we had difficulties fitting the reaction rates to fluorescent signal for toehold lengths 1 and 2, presumably due to the weak signal that has not saturated even over the course of the duration of the experimental measurements. The experimental measurements are shown in Fig. S5.

We next compared the dependence of the TMSD kinetics on the toehold placement at either 5′ or 3′ end of the substrate. The rate dependence on whether the toehold is at the 5’ or 3’ end have not been reported in a systematic study for RNA strand displacement, however the coarse-grained simulations of TMSD for RNA observed the speed-up of the reaction if toehold is placed at the 5′of the substrate [30]. This phenomena was observed due to the additional cross-stacking interaction between the invader and 5’ end toehold substrate strand due to the A-form helix structure of the RNA duplex, which provides extra stabilization as opposed to when the the invader binds to 3′ toehold. The simulations predict that strand displacement is faster by about factor 2 to 10 for shorter toeholds until the rate of TMSD is saturated (which is at about 6 base pairs for averaged strength sequences) [30].

We measured the rate of TMSD for toeholds of length 1, 3 and 6 nts placed at 3′ end, which were designed so that the free energy of invader binding to them would be the same as the free energy of binding to toehold of the same length at 5′ end (see Methods and Supp. Mat.). The fits of the rate of displacement for toehold lengths 3 nt show that the fitted TMSD rate *k*_eff_ placed at the 3′ end is comparable to the 5′ end and the fitted effective rate *k*_eff_ is comparable for both 3′> and 5′ ends. For the toehold length 6 nts, the toehold placed at 5’ end had approximately twenty times faster rate of TMSD than the one placed at the 3’ end. We were unable to fit the rates for toeholds of length 1 nt, as the signal was very weak over the duration of the experiment. These results are in contrast with the simulations, where the rates of displacament were faster for the 5’ end placement for 3nt length by about a factor of 2, and were comparable for 6nt length. It is possible that there are additional sequence-dependent effects that can play role in the TMSD kinetics.

### B. Mismatch effects

We further study the effects of mismatches between the invader RNA strand and the RNA substrate. For each system studied, we introduce a single mismatch in the RNA invader which cannot form Watson-Crick nor wobble complementary with the corresponding base on the substrate. For all the experiments with the mismatches, we consider substrate with toehold length 6 nt placed at the 5′ end. We only introduce a single mismatch in each experimentally studied system. We consider 6 different mismatch positions in the invader, as shown schematically in Fig. 2a. The mismatches are present either in the toehold binding region (P25, P23 or P21), in the first base pair next to the toehold end (P20), in the middle of invader (P10) and three bases from the end (P3). The experimental measurements are shown in Fig. 3 along with the fitted rates in Table I and in Fig. 4.

**FIG. 4.**
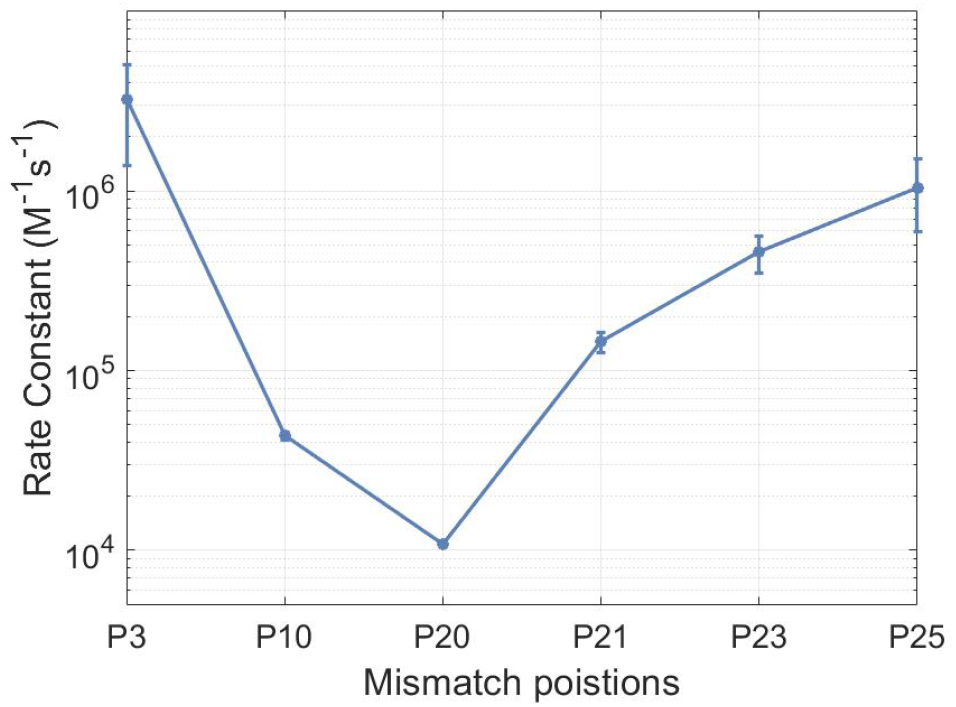
The fitted rate constants of reactions with different mismatches.

We observe that all considered mismatch positions lead to a slow-down of the reaction. However, the effects of the mismatch depend on the position with respect to the interface where the toehold region ends and the incumbent region begins: position P21 of the invader is the last base of the toehold region and P21 is the first base which displaces the incumbent. We observe that the largest slowdown (by about a factor of 134) is for the mismatch at P20 (Fig. 4), and the smallest effect for the mismatch at the end of furthest point away from the toehold (P3). For the mismatches in the toehold region (P25-P21), we find that the closer the mismatch is to the interface, the bigger the slowdown (by up to a factor of 7 difference between the furthest and closest points). Finally, for the mismatches in the incumbent displacement region, mismatches at P3 and P10, the effect of slowdown of the displacement reaction by a factor of 36 is observed for mismatch at P10 which is closer to the toehold while for mismatch at P3 no significant slowdown effect is identified.

The observed slowdown induced by mismatches at different positions for RNA TMSD is hence compatible with the kinetics of TMSD of DNA strands with mismatches that was previously studied computationally and experimentally [26, 27, 34]. The kinetics of the TMSD can be approximately modeled as a stochastic Markovian process consisting of transitions between states characterized by the number of bonds between respective strands. Introducing a mismatch between the invader and the substrate creates a free-energy barrier which increases the time it takes the invader to displace the substrate. The kinetics of the strand displacement is more affected by the mismatches closer to the toehold interface, as it increases the likelihood of the invader detaching from the toehold before successfully displacing the incumbent. Mismatches closer to the interface increase the barrier to the displacement initiation even more, thus having larger effect. Mismatches close to the end (P3) have smaller effect on the kinetic rate, as it is also possible for the substrate strand to spontaneously detach while it has only three remaining base pairs.

### C. RNA:DNA hybrid strand displacement

We next compare the hybrid TMSD systems where either DNA invades RNA duplex (DNAtoRNA) or RNA invades a DNA duplex (RNAtoDNA). For the studied toehold lengths (1, 3 and 6 nt), we also compare the hybrid TMSD systems with experiments where RNA invades RNA duplex (RNAtoRNA) and where DNA invades DNA duplex (DNAtoDNA). For the corresponding experiments at a given toehold length, we use the same nucleic acids sequences for invader, substrate and incumbent strands. For all the studied hybrid systems and DNAtoDNA systems, the toehold is placed at the 5’ end of the substrate. The fitted TMSD rates are shown in Table I with the corresponding curves shown in Fig. 3d. We used the same sequence (with substituting U for T in the case of RNA), as listed in the Supplementary Material (Tables S1-S4).

For the longest toehold considered (6 nts), we observe that TMSD for DNA invading DNA duplex the fitted TMSD rates *k*_eff_ sorted from the fastest to slowest are: RNAtoRNA, DNAtoDNA, RNAtoDNA with fitted rates log_10_ *k*_eff_ : 6.3, 6.0, and 5.7 respectively with DNAtoRNA not fittable but with clearly slowest rate from comparing them in Fig. 3c. For the shorter toehold length of 3 nt, the RNAtoRNA appears to be the fastest based on the analysis of the fluorescent signal (Fig. 3), followed by DNAtoDNA and then RNAtoDNA. We did not study DNAtoRNA for 3nt toehold. We note that RNAtoDNA and DNAtoDNA reactions could not have been reliably fit with with our model (see Table I), presumably since the reaction was too slow and did not reach equilibrium with the time of the experiment.

We rationalize the observed behavior of hybrid strand displacement through free-energy landscapes and known thermodynamic stabilities of duplexes of DNA, RNA, and RNA:DNA hybrids respectively [35–37]. The estimated free-energy profiles, based on intuitive energy landscape from Ref. [24] are shown in Figure S8 for the TMSD with 6nt and 3nt toehold. For the same sequence, RNA:RNA toehold binding has the largest stability, followed by RNA:DNA and DNA:DNA respectively. Such free-energy landscapes have been used in prior studies for DNA [38]. Once the invader is bound to the toehold, it can either dissociate (overcoming the barrier given by binding to the toehold) or it can displace the incumbent. After overcoming the initial barrier to the displacement (about 2 *k*_B_*T*), the invader replaces the incumbent through branch migration process. For RNAtoRNA or DNAtoDNA TMSD systems, the newly created base pairs have identical stability, while for RNAtoDNA, the new bases are on average more stable than the original ones, leading to a downhill free-energy landscape. For DNAtoRNA, the newly created base pairs are less stable, so the branch migration part of TMSD corresponds to an uphill free-energy landscape. We note, however, that DNAtoDNA reaction appears slightly faster than RNAtoDNA for 6nt toehold system, despite the fact that the free-energy landscape is downhill for the RNAtoDNA system due to the higher stability of the newly created base pairs. Since RNA:DNA hybrid duplex is known to prefer A-form helix, as opposed to the B-form typical for DNA duplexes, we hypothesize that the rearrangement from B-form helix into A-form might also affect the displacement rate.

The higher stability of RNA:RNA toehold than RNA:DNA toehold likely contributes to the slightly faster rate for RNAtoRNA 6 nt toehold than for the RNAtoDNA and DNAtoDNA systems. For the sequences used for toehold of 3 nts length, the stability of RNA:RNA and RNA:DNA toehold is comparable, and hence the RNAtoDNA is expected to proceed faster due to the higher stability of newly formed RNA base pairs in the branch migration region.

## IV. CONCLUSION

In a series of experiments, we found the RNA TMSD to show similar phenomena observed in DNA in terms of the toehold length effects and mismatches between the invader and the substrate. Through the systematic study of RNA and RNA:DNA hybrid strand displacement, we discovered that (i) Increasing the toehold length has a significant boost on strand displacement reaction rate, resulting in exponential speed-up of the displacement reaction until saturation speed is reached, consistent with phenomena previously observed for DNA TMSD [6]; (ii) Placement of the RNA toehold at 5’ end of substrate can accelerate the reaction for 6nt toehold; (iii) Base mismatches on invader generally slows down the reaction, and the closer they are to to the interface of toehold and branch migration range the bigger the effect; (iv) For toehold of length 6nt, RNA invading RNA duplex is faster than RNA invading DNA duplex. For toehold of length 3 nt, we found RNA invading DNA TMSD to be faster than RNA invading RNA.

As the emerging field of RNA nanotechnology evolves and the *in vivo* applications, understanding the kinetics of RNA strand displacement is of increasing importance. Furthermore, hybrid displacement systems that interface DNA with RNA are further of interest for DNA nanotechnology interfacing with biological systems. Finally, strand displacement reactions for RNA and hybrid systems can also play role not only in designed synthetic systems, but also in biological processes. Our results provide first systematic study of RNA and DNA:RNA hybrid displacement kinetics and will help in understanding the phenomena and building stochastic models of TMSD reactions for RNA.

## V. ACKNOWLEDGEMENTS

The authors thank Shuoxing Jiang, Erik Poppleton, Michael Matthies, Xu Zhou and Lan Liu for helpful discussions.

## References

[1] S. Li, Q. Jiang, S. Liu, Y. Zhang, Y. Tian, C. Song, J. Wang, Y. Zou, G. J. Anderson, J.-Y. Han, et al. A DNA nanorobot functions as a cancer therapeutic in response to a molecular trigger in vivo. Nature biotechnology, 36(3):258, 2018.

[2] X. Qi, X. Liu, L. Matiski, R. Rodriguez Del Villar, T. Yip, F. Zhang, S. Sokalingam, S. Jiang, L. Liu, H. Yan, et al. RNA Origami Nanostructures for Potent and Safe Anticancer Immunotherapy. ACS nano, 14(4):4727–4740, 2020.

[3] K. Pardee, A. A. Green, M. K. Takahashi, D. Braff, G. Lambert, J. W. Lee, T. Ferrante, D. Ma, N. Donghia, M. Fan, et al. Rapid, low-cost detection of Zika virus using programmable biomolecular components. Cell, 165(5):1255–1266, 2016.

[4] P. Guo. The emerging field of RNA nanotechnology. Nature nanotechnology, 5(12):833, 2010.

[5] F. C. Simmel, B. Yurke, and H. R. Singh. Principles and applications of nucleic acid strand displacement reactions. Chemical reviews, 119(10):6326–6369, 2019.

[6] D. Y. Zhang and E. Winfree. Control of DNA strand displacement kinetics using toehold exchange. Journal of the American Chemical Society, 131(47):17303–17314, 2009.

[7] B. Yurke, A. J. Turberfield, A. P. Mills Jr, F. C. Simmel, and J. L. Neumann. A DNA-fuelled molecular machine made of DNA. Nature, 406(6796):605, 2000.

[8] B. Wei, L. L. Ong, J. Chen, A. S. Jaffe, and P. Yin. Complex reconfiguration of DNA nanostructures. Angewandte Chemie, 126(29):7605–7609, 2014.

[9] G. Grossi, M. D. E. Jepsen, J. Kjems, and E. S. Andersen. Control of enzyme reactions by a reconfigurable DNA nanovault. Nature communications, 8(1):992, 2017.

[10] S. F. Wickham, M. Endo, Y. Katsuda, K. Hidaka, J. Bath, H. Sugiyama, and A. J. Turberfield. Direct observation of stepwise movement of a synthetic molecular transporter. Nature nanotechnology, 6(3):166–169, 2011.

[11] L. Qian, E. Winfree, and J. Bruck. Neural network computation with DNA strand displacement cascades. Nature, 475(7356):368–372, 2011.

[12] D. Wilhelm, J. Bruck, and L. Qian. Probabilistic switching circuits in DNA. Proceedings of the National Academy of Sciences, 115(5):903–908, 2018.

[13] L. Qian and E. Winfree. Scaling up digital circuit computation with DNA strand displacement cascades. Science, 332(6034):1196–1201, 2011.

[14] F. Hong and P. Šulc. An emergent understanding of strand displacement in RNA biology. Journal of structural biology, 207(3):241–249, 2019.

[15] M. Y. Angela, P. M. Gasper, L. Cheng, L. B. Lai, S. Kaur, V. Gopalan, A. A. Chen, and J. B. Lucks. Computationally reconstructing cotranscriptional RNA folding from experimental data reveals rearrangement of non-native folding intermediates. Molecular Cell, 2021.

[16] A. A. Green, P. A. Silver, J. J. Collins, and P. Yin. Toehold switches: de-novo-designed regulators of gene expression. Cell, 159(4):925–939, 2014.

[17] M. Klein, B. Eslami-Mossallam, D. G. Arroyo, and M. Depken. Hybridization kinetics explains CRISPR-Cas off-targeting rules. Cell reports, 22(6):1413–1423, 2018.

[18] C. Anders, O. Niewoehner, A. Duerst, and M. Jinek. Structural basis of PAM-dependent target DNA recognition by the Cas9 endonuclease. Nature, 513(7519):569, 2014.

[19] E. A. Josephs, D. D. Kocak, C. J. Fitzgibbon, J. McMenemy, C. A. Gersbach, and P. E. Marszalek. Structure and specificity of the RNA-guided endonuclease Cas9 during DNA interrogation, target binding and cleavage. Nucleic acids research, 43(18):8924–8941, 2015.

[20] I. Strohkendl, F. A. Saifuddin, J. R. Rybarski, I. J. Finkelstein, and R. Russell. Kinetic basis for DNA target specificity of CRISPR-Cas12a. Molecular cell, 71(5):816–824, 2018.

[21] C. M. Radding, K. L. Beattie, W. K. Holloman, and R. C. Wiegand. Uptake of homologous single-stranded fragments by superhelical DNA: IV. Branch migration. Journal of Molecular Biology, 116(4):825–839, 1977.

[22] C. Green and C. Tibbetts. Reassociation rate limited displacement of DNA strands by branch migration. Nucleic Acids Research, 9(8):1905–1918, 04 1981.

[23] B. Yurke and A. P. Mills. Using DNA to power nanostructures. Genetic Programming and Evolvable Machines, 4(2):111–122, 2003.

[24] N. Srinivas, T. E. Ouldridge, P. Šulc, J. M. Schaeffer, B. Yurke, A. A. Louis, J. P. Doye, and E. Winfree. On the biophysics and kinetics of toehold-mediated DNA strand displacement. Nucleic acids research, of Sciences, 115(5):903–908, 2018.41(22):10641–10658, 2013.

[25] T. E. Ouldridge, A. A. Louis, and J. P. Doye. Structural, mechanical, and thermodynamic properties of a coarsegrained DNA model. The Journal of chemical physics, 134(8):02B627, 2011.

[26] R. R. Machinek, T. E. Ouldridge, N. E. Haley, J. Bath, and A. J. Turberfield. Programmable energy landscapes for kinetic control of DNA strand displacement. Nature communications, 5(1):1–9, 2014.

[27] N. E. Haley, T. E. Ouldridge, A. Geraldini, A. A. Louis, J. Bath, and A. J. Turberfield. Rational design of hidden thermodynamic driving through DNA mismatch repair. bioRxiv, page 426668, 2018.

[28] D. B. Broadwater Jr and H. D. Kim. The effect of basepair mismatch on DNA strand displacement. Biophysical journal, 110(7):1476–1484, 2016.

[29] P. Šulc, F. Romano, T. E. Ouldridge, J. P. Doye, and A. A. Louis. A nucleotide-level coarse-grained model of RNA. The Journal of chemical physics, 140(23):06B614 1, 2014.

[30] P. Šulc, T. E. Ouldridge, F. Romano, J. P. Doye, and A. A. Louis. Modelling toehold-mediated RNA strand displacement. Biophysical journal, 108(5):1238–1247, 2015.

[31] D. Broadwater Jr, A. W. Cook, and H. D. Kim. First passage time study of DNA strand displacement. arXiv preprint 2005.10925, 2020.

[32] J. N. Zadeh, C. D. Steenberg, J. S. Bois, B. R. Wolfe, M. B. Pierce, A. R. Khan, R. M. Dirks, and N. A. Pierce. NUPACK: Analysis and design of nucleic acid systems. Journal of computational chemistry, 32(1):170–173, 2011.

[33] M.-X. Li, C.-H. Xu, N. Zhang, G.-S. Qian, W. Zhao, J.-J. Xu, and H.-Y. Chen. Exploration of the kinetics of toehold-mediated strand displacement via plasmon rulers. ACS nano, 12(4):3341–3350, 2018.

[34] P. Irmisch, T. E. Ouldridge, and R. Seidel. Modelling DNA-strand displacement reactions in the presence of base-pair mismatches. Journal of the American Chemical Society, 2020.

[35] J. SantaLucia. A unified view of polymer, dumbbell, and oligonucleotide DNA nearest-neighbor thermodynamics. Proceedings of the National Academy of Sciences, 95(4):1460–1465, 1998.

[36] N. Sugimoto, S.-i. Nakano, M. Katoh, A. Matsumura, H. Nakamuta, T. Ohmichi, M. Yoneyama, and M. Sasaki. Thermodynamic parameters to predict stability of RNA/DNA hybrid duplexes. Biochemistry, 34(35):11211–11216, 1995.

[37] M. J. Serra and D. H. Turner. [11] Predicting thermodynamic properties of RNA. In Methods in enzymology, volume 259, pages 242–261. Elsevier, 1995.

[38] T. E. Ouldridge, P. Šulc, F. Romano, J. P. Doye, and A. A. Louis. DNA hybridization kinetics: zippering, internal displacement and sequence dependence. Nucleic acids research, 41(19):8886–8895, 2013.

