## Supplementary Information for "Kinetics of RNA and RNA:DNA hybrid strand displacement"

### SI. SEQUENCE DESIGN

The sequences were designed and analyzed using NUPACK [1]. Sequences of all the strands used in this study are listed in following tables. Each strand was named by the experiment they were used. All the sequences are listed with an order of 5' to 3' order. For all invader strands for mismatch study, the position that designed to be mismatching with the substrate strand is labeled in a fashion as P3, P10 etc. (the third nt counted from 5' end).

*

| Strand | Sequence |
|---|---|
| subom5_strand | GACCAGGAGCAGCGAAGAUAGAUAUG |
| incumbent5_strand | UCCUCUGUCUCUAUUCAUAUCUAUCUUCGCUGCUC |
| invad5_strand | CAUAUCUAUCUUCGCUGCUCCUGGUC |
| mut_invad5_p3 | CACAUCUAUCUUCGCUGCUCCUGGUC |
| mut_invad5_p10 | CAUAUCUAUAUUCGCUGCUCCUGGUC |
| mut_invad5_p20 | CAUAUCUAUCUUCGCUGCUACUGGUC |
| mut_invad5_p21 | CAUAUCUAUCUUCGCUGCUCAUGGUC |
| mut_invad5_p23 | CAUAUCUAUCUUCGCUGCUCCUAGUC |
| mut_invad5_p25 | CAUAUCUAUCUUCGCUGCUCCUGGAC |
| sub5_strand_t1 | GGAGCAGCGAAGAUAGAUAUG |
| inv5_strand_t1 | CAUAUCUAUCUUCGCUGCUCC |
| sub5_strand_t2 | GGGAGCAGCGAAGAUAGAUAUG |
| inv5_strand_t2 | CAUAUCUAUCUUCGCUGCUCCC |
| sub5_strand_t3 | GAGGAGCAGCGAAGAUAGAUAUG |
| inv5_strand_t3 | CAUAUCUAUCUUCGCUGCUCCUC |
| sub5_strand_t4 | GAAGGAGCAGCGAAGAUAGAUAUG |
| inv5_strand_t4 | CAUAUCUAUCUUCGCUGCUCCUUC |
| sub5_strand_t5 | GACAGGAGCAGCGAAGAUAGAUAUG |
| inv5_strand_t5 | CAUAUCUAUCUUCGCUGCUCCUGUC |

TABLE S1. The sequences of all strands used to explore the effect of different toehold length and mismatch positions to reaction kinetics. Toehold is located at the 5' end for all strands listed in this table.

| Strand | Sequence |
|---|---|
| subom3_strand | GAGCAGCGAAGAUAGAUAUGGACCAG |
| incumbent3_strand | CAUAUCUAUCUUCGCUGCUCUUACAUUCAUCAGUC |
| invader3_strand | CUGGUCCAUAUCUAUCUUCGCUGCUC |
| sub3_strand_t1 | GAGCAGCGAAGAUAGAUAUGG |
| inv3_strand_t1 | CCAUAUCUAUCUUCGCUGCUC |
| sub3_strand_t2 | GAGCAGCGAAGAUAGAUAUGGG |
| inv3_strand_t2 | CCCAUAUCUAUCUUCGCUGCUC |
| sub3_strand_t3 | GAGCAGCGAAGAUAGAUAUGGAG |
| inv3_strand_t3 | CUCCAUAUCUAUCUUCGCUGCUC |
| sub3_strand_t4 | GAGCAGCGAAGAUAGAUAUGGAAG |
| inv3_strand_t4 | CUUCCAUAUCUAUCUUCGCUGCUC |
| sub3_strand_t5 | GAGCAGCGAAGAUAGAUAUGGACAG |
| inv3_strand_t5 | CUGUCCAUAUCUAUCUUCGCUGCUC |

TABLE S2. The sequence of all strands used to explore the effect of toehold locations to reaction kinetics. Toehold is located at the 3' end for all strands listed in this table.

| Strand | Sequence |
|---|---|
| substrate5_strand_DNA | GACCAGGAGCAGCGAAGATAGATATG |
| incumbent5_strand_DNA | TCCTCTGTCTCTATTCATATCTATCTTCGCTGCTC |
| invad5_strand_DNA | CATATCTATCTTCGCTGCTCCTGGTC |
| mut_invad5_p3_DNA | CACATCTATCTTCGCTGCTCCTGGTC |
| mut_invad5_p10_DNA | CATATCTATATTCGCTGCTCCTGGTC |
| mut_invad5_p20_DNA | CATATCTATCTTCGCTGCTACTGGTC |
| mut_invad5_p21_DNA | CATATCTATCTTCGCTGCTCATGGTC |
| mut_invad5_p23_DNA | CATATCTATCTTCGCTGCTCCTAGTC |
| mut_invad5_p25_DNA | CATATCTATCTTCGCTGCTCCTGGAC |
| sub5_strand_t1_DNA | GGAGCAGCGAAGATAGATATG |
| inv5_strand_t1_DNA | CATATCTATCTTCGCTGCTCC |
| sub5_strand_t2_DNA | GGGAGCAGCGAAGATAGATATG |
| inv5_strand_t2_DNA | CATATCTATCTTCGCTGCTCCC |
| sub5_strand_t3_DNA | GAGGAGCAGCGAAGATAGATATG |
| inv5_strand_t3_DNA | CATATCTATCTTCGCTGCTCCTC |
| sub5_strand_t4_DNA | GAAGGAGCAGCGAAGATAGATATG |
| inv5_strand_t4_DNA | CATATCTATCTTCGCTGCTCCTTC |
| sub5_strand_t5_DNA | GACAGGAGCAGCGAAGATAGATATG |
| inv5_strand_t5_DNA | CATATCTATCTTCGCTGCTCCTGTC |

TABLE S3. The sequence of all strands of Table S1 in DNA used for the study of DNA:DNA and RNA:DNA hybrid strand displacement.

| Strand | Sequence |
|---|---|
| reporter_sub5_strand_TAM | GATATGAATAGAGACAGAGGA/36-TAMTSp/ |
| reporter_up5_strand_FAM | /56-FAM/TCCTCTGTCTCTATT |
| reporter_sub3_strand | /56-TAMN/GACTGATGAATGTAAGAGCAG |
| reporter_up3_strand | TTACATTCATCAGTC/36-FAM/ |

TABLE S4. All strands used for fluorescence readout including the cases for 5' toehold and 3' toehold. TAMTSp and TAMN are two types of TAMRA fluorophores used in this study.

### SII. RNA SECONDARY STRUCTURE AND DEGRADATION

The chemical instability of RNA comes from the extra 2' hydroxyl group compared to DNA. While it makes RNA chemically reactive and easy to modify at 2' position, it also makes RNA susceptible degradation in the environment send self-cleavage. Besides the problem of degradation in a quantitative experiment, the non-canonical base pairing (G-U) also offers more possibilities to form secondary structure than the same sequence would have in DNA. In the experiment we seek to avoid the RNA degradation and secondary structure formation, as discussed below.

#### A. NUPACK predictions on secondary structures

We designed the sequences to minimize the undesired secondary structure for most of the strands used in this study. While for some strands whose secondary structure is confirmed by NUPACK (i.e. Mismatch at P25 in Fig S2), in the analysis they are excluded for the accuracy. In NUPACK predictions on secondary structures from Figure S1-S3, parameters are set to be 25°C, 60 nM, maximum complex size 4 strands. For most RNA strands, even with certain extent of secondary structure, the equilibrium probability is limited and for most substrate strands that could form secondary structure, annealing with incumbent strand could solve the problem.

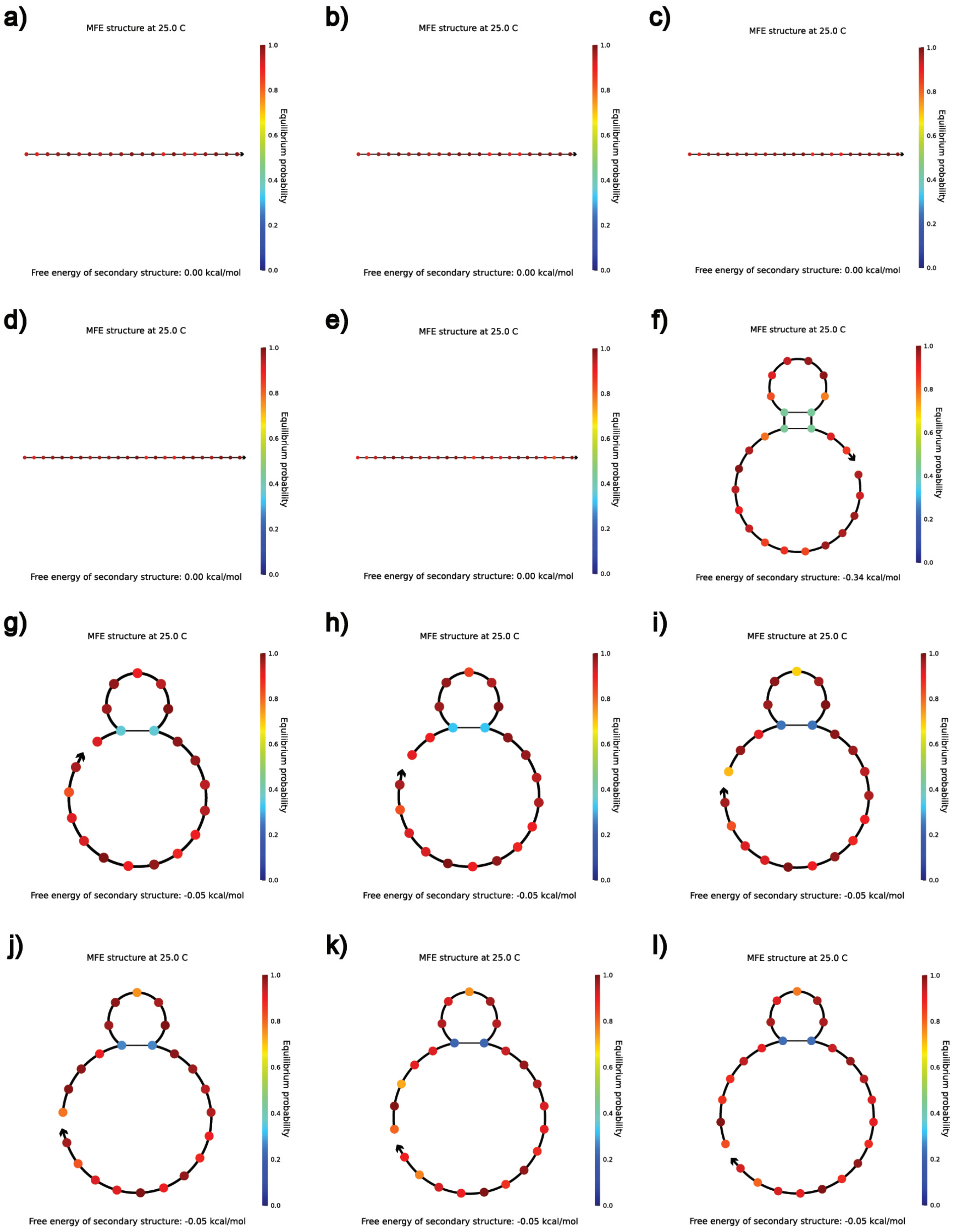


FIG. S1. NUPACK predictions on 5' toehold RNA invader and substrate strands with toehold length varied from 1nt to 6nt. Invader with toehold being a) 1nt b)2nt c)3nt d)4nt e)5nt f)6nt. Substrate with toehold being g)1nt h)2nt i)3nt j)4nt k)5nt l)6nt.

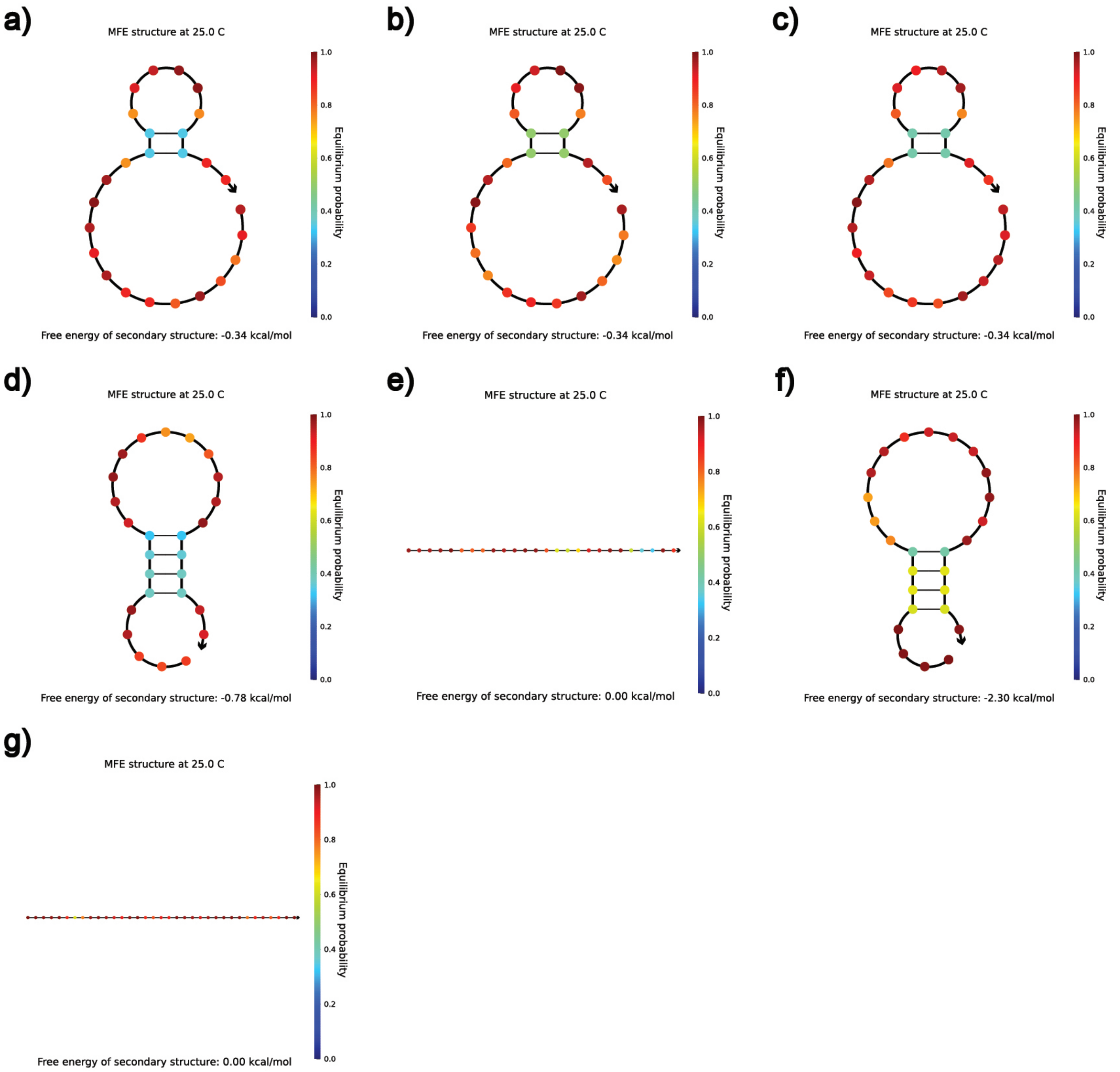


FIG. S2. NUPACK predictions on RNA invader strands with mismatch positions and incumbent strand. Counting from the 5' end, mismatch is placed at a) position 3 b) position 10 c) position 20 d)position 21 e) position 23 f) position 25. g) Incumbent strands with toehold placed at 5' end.

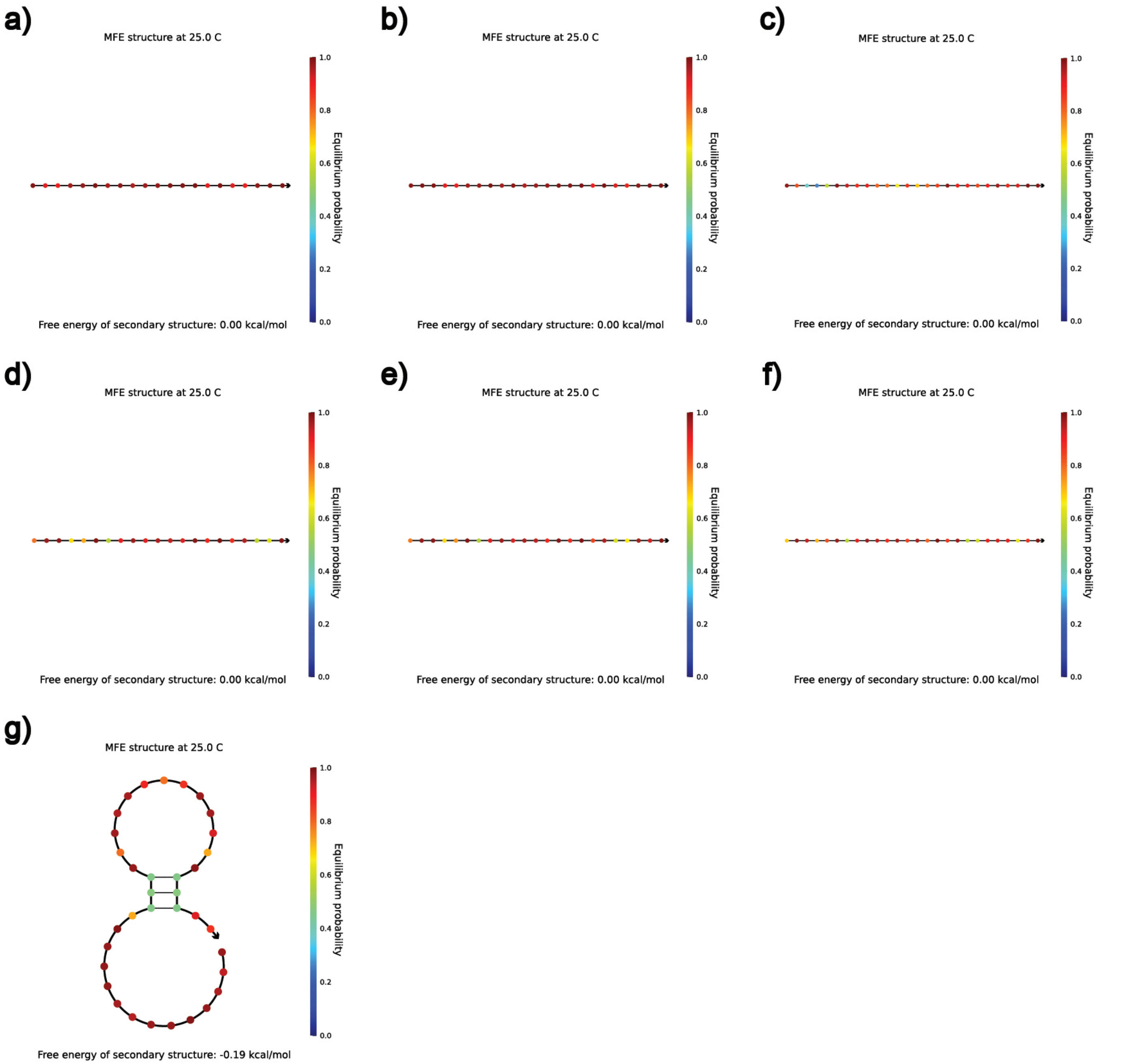


FIG. S3. NUPACK predictions on 3' toehold RNA invader,substrate and incumbent strands with different toehold lengths. Invader with toehold being a) 1nt b) 3nt c) 6nt. Substrate with toehold being d) 1nt e) 3nt f) 6nt. g) Incumbent strands with toehold placed at 3' end.

#### B. Gel electrophoresis

To further verify 1) whether RNA strands form secondary structure in experiment 2) whether RNA degrades during the purification and dilution, we run native gels to test RNA strands subjected to fluorescence measurement due to the buffer requirement. In Figure S4, four gel pictures are shown for different experiment series. While the invader of 5' toehold RNA strands with length ranging from 1nt to 6nt exhibit no signs of secondary structures or degradation, the 5nt gate and 2nt gate behaves out of our expectation. It is clear that the upper smear of 5nt gate is due to the formation of secondary structure and it is confirmed by the kinetics curves (Figure S4) as it is unusually slow for 5nt case and we hence did not use the 5nt toehold design for comparison with other TMSDs in this work, as the kinetics is affected by secondary structure formation. As for the double band of the 2nt toehold design, we assume it can be ascribed to the inexact stochiometry.

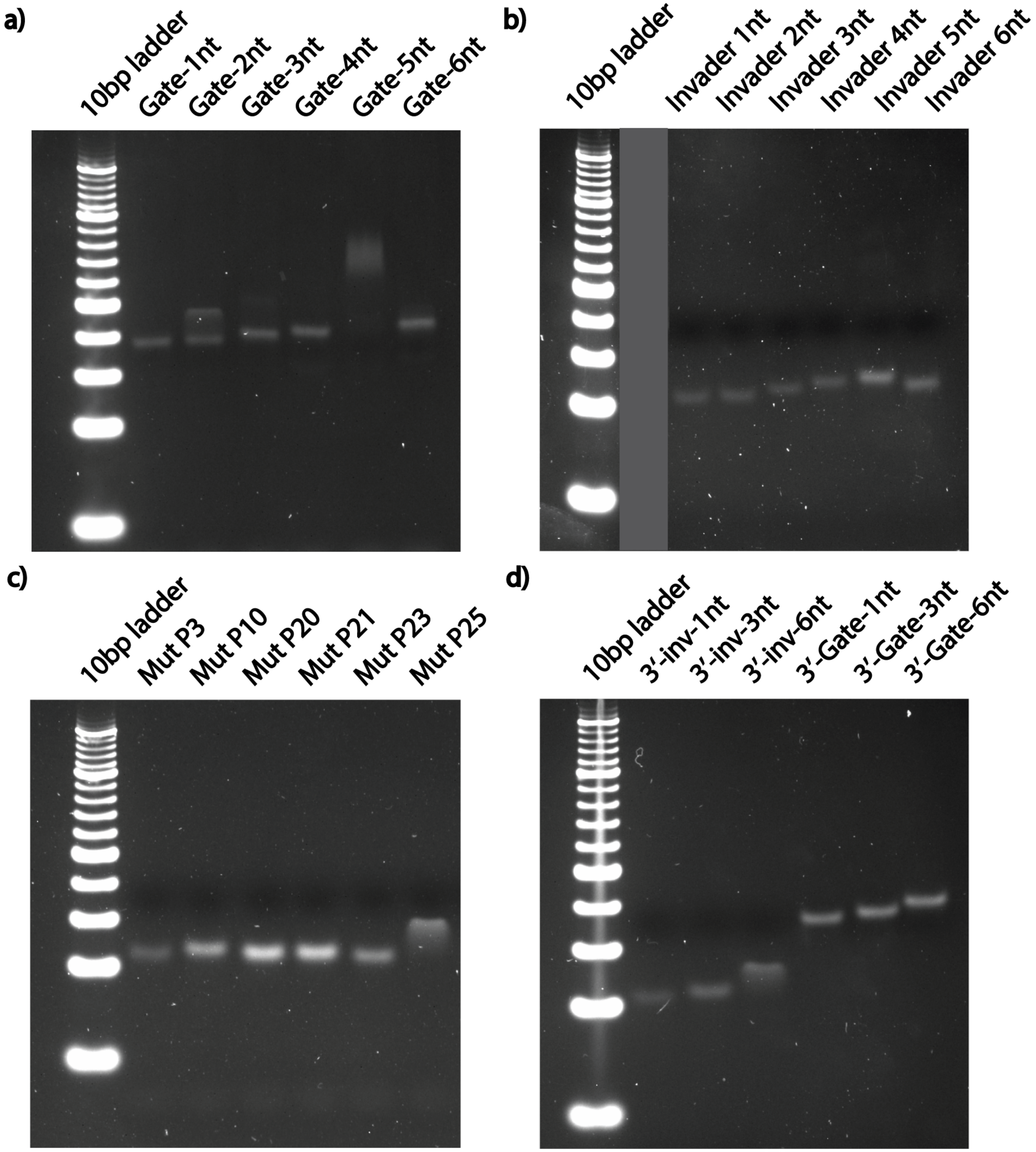


FIG. S4. Gel electrophoresis on RNA strands. The first part is native gel on 5' toehold RNA series: a) gates annealed with incumbent and substrate strand whose toehold length varies from 1nt to 6nt. b) Single stranded invader with toehold length from 1nt to 6nt. The mismatch series is shown in c) with invader strand positioned mismatching nt at different places. In d), the 3' toehold RNA series with toehold being 1, 3, 6 nt are shown.

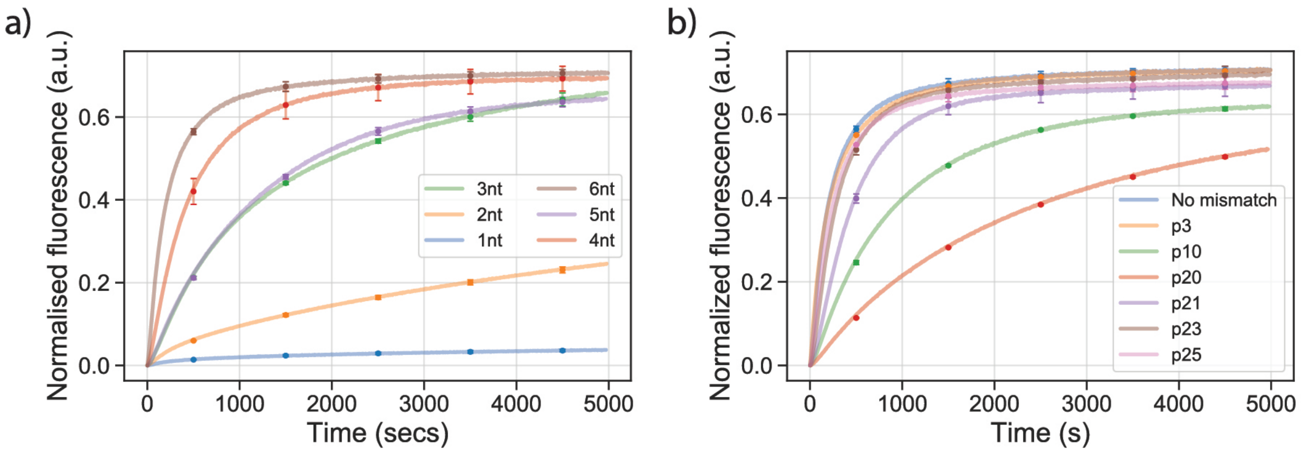


FIG. S5. Full curves of the a) 5' toehold and b)mismatch study experiments including the one with secondary structure.

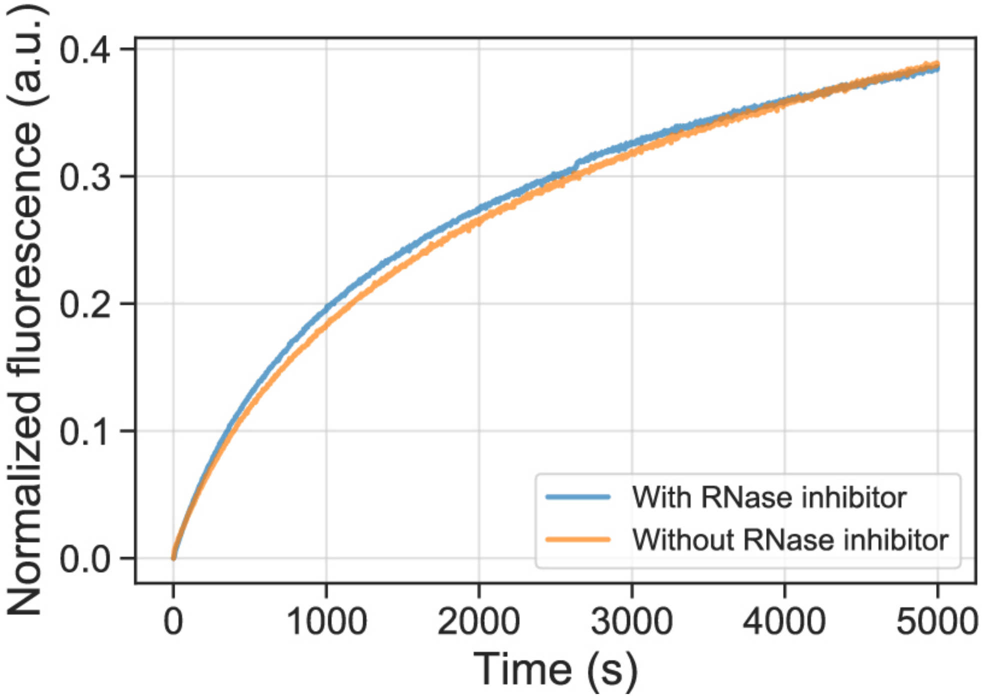


FIG. S6. Kinetic curves of a chosen reaction (toehold at 3' with 3nt length) with or without the addition of the RNase inhibitor during the fluorescent measurement.

#### C. Fluorescent measurement control

To evaluate the influence of RNA degradation during the fluorescent measurement, we chose one RNA TMSD reaction with 3nt toehold located at 3'. We ran the same reaction with and without RNase inhibitor added (Figure S6). The concentration of the RNase inhibitor is maintained as 1 unit/ul.

From comparison of the curves, we observe no significant difference and hence conclude that over the course of the experiment, there is no significant degradation in our experimental setup. We hence carried out the experiments without the RNase inhibitor.

### SIII. CURVE FITTING

Two methods are adopted in this study to fit kinetic curves, namely method I adapting the strategies reported by Zhang and Winfree [2] and method II adapted from Machinek et al[3] with assumption that the reporter reaction is much faster than the original strand displacement. For some reactions (1 nt toehold length or 5nt toehold with secondary structure formation) the fitting quality (R scores of the fitting curve) is not as good as we expected and for some reactions (DNAtoDNA 3nt toehold or RNAtoRNA 2nt toehold) the rate constant predicted by the model is not plausible, as it would correspond to unreasonable values of $\alpha$, so even though the curve can be fitted with the model, the resulting rate is unphysical. We hypothesize that this might be due to the stochiometric ratio of the invader and substrate along with the reporter: while such experimental set-up clearly differentiates the fast and slow reaction kinetics with different screening parameters, i.e, toehold length, it makes the curve corresponding to the slow reactions recorded within 5000 seconds not able to reflect the kinetic properties enough for the fitting. We tried to elongate the observing time to 15000 seconds for the slow reactions and the signal remain growing in a slow and almost linear fashion with the fact of reaching a plateau no foreseeable (data not shown). In this regard we hypothesize that using high excess of invader/substrate concentration might lead to a reaction that would be more easily fitable. Nonetheless, the trend still holds as after excluding 1nt and 5nt the monotonically increasing of rate constants is observed for 5' toehold case and similarly the expected or explainable trend is found for other experimental series. We further discuss the fitting methods below.

#### A. Method I

The reporter characterisation reactions are represented as:

$$\text{Incumbent} + \text{Reporter} \xrightarrow{k_{\text{rep}}} \text{Signal} + \text{WasteT}$$

Where we label the species as: Incumbent (I), Reporter (R), WasteT (W), Signal (S). We use the following set of ODEs to capture the reaction kinetics:

$$\frac{\mathrm{d(S)}}{\mathrm{d(t)}} = \frac{\mathrm{d(W)}}{\mathrm{d(t)}} = -\frac{\mathrm{d(R)}}{\mathrm{d(t)}} = \frac{\mathrm{d(I)}}{\mathrm{d(t)}} = \mathrm{k_{eff}[I][R]}$$

The fitting code is available for download at https://github.com/sulcgroup/TMSD_fitting.git. Briefly, we defined a function TMSD_system, which defined the relevant set of ODEs, with function inputs of time, t, the initial reactant concentrations and $k_{\text{rep}}$ and function outputs of $\frac{\mathrm{d(S)}}{\mathrm{d(t)}}$, $\frac{\mathrm{d(R)}}{\mathrm{d(t)}}$, $\frac{\mathrm{d(I)}}{\mathrm{d(t)}}$ and $\frac{\mathrm{d(W)}}{\mathrm{d(t)}}$. In order to identify best-fit values for $\alpha$ and $k_{\text{rep}}$. scipy.optimize.curve_fit takes the integral of $\frac{\mathrm{d(S)}}{\mathrm{d(t)}}$, t and the initial concentration of F as inputs. We provided upper and lower bounds for the fitted rate constants and scaling factor within the curve_fit function. For $k_{\text{rep}}$ and $\alpha$, we provided bounds of 1 to 7, and 0 to 1, respectively. We also provided an initial estimate of $k_{\text{rep}}$ and values of 6 and 0.8, respectively. However, we noted that the initial estimate did not seem to influence the fitted values greatly. scipy.optimize.curve_fit uses a non-linear least squares approach to the fit the integral of $\frac{\mathrm{d(S)}}{\mathrm{d(t)}}$ to the experimental data and provide estimates for $k_{\text{rep}}$ and $\alpha$ (Table S5).

For the complete 2-step strand displacement reactions, we proceed analogously. We have the following definitions and ODEs:

$$\text{Invader} + \text{Gate} \xrightarrow{k_{\text{eff}}} \text{Incumbent} + \text{WasteB}$$
$$\text{Incumbent} + \text{Reporter} \xrightarrow{k_{\text{rep}}} \text{Signal} + \text{WasteT}$$

$$\frac{d(V)}{d(t)} = -\frac{d(G)}{d(t)} = k_{eff}[V][G]$$

$$\frac{d(I)}{d(t)} = k_{eff}[V][G] - k_{rep}[I][R]$$

$$\frac{d(S)}{d(t)} = -\frac{d(R)}{d(t)} = k_{rep}[I][R]$$

We implemented a similar program to fit $k_{\text{eff}}$ with the fitted $k_{\text{rep}}$ from above fitting process.

In Figure S7, it can be clearly seen that that the model fails to fit 1nt toehold case. We think this is due to the fact that for the 1nt case the equilibrium is not reached within 5000s. That is a universal problem for reactions with slow reaction rate. However, increasing the data collection time further would increase the difficulty of carrying out the experiment would be to increase data collection time with longer time, which would increase the difficulty of carrying out the experiment, as well as increase the risk of degradation of RNA.

We further experienced issues with fitting the slower reactions, where we were able to fit the curves, but the resulting rates $k_{\text{eff}}$ were unreasonable, sometimes the resulting fit had very high values of $k_{\text{eff}}$ compensated by unreasonably low value of $\alpha$. As shown in Table S6, slow reactions are predicted to have relatively high rate constants and even exhibiting the reversed trend revealed by the direct comparison of kinetic curves.

### B. Method II

We also adapted another fitting method descried by Machinek et al [3] which assumes the reporter reaction is much faster than the displacement itself and therefore it should be particularly suitable for slow displacement reaction like short toehold length cases. The equation is solved from the relation of all species and represented as:

$$[R] = \frac{I_0{}^2kt}{1+I_0kt}$$

The equation to be fitted in here is a rough approximation of the actual kinetics. The prerequisite is that reporter is displaced with a much faster rate compared to the release of incumbent and for reaction with fast rates the prediction on rate constant can be expected to be inaccurate. For fast reactions, the predicted rate constants turns out to be underestimated when compared to the result of Method I, which is expected due to to the assumptions employed (Figure S7).

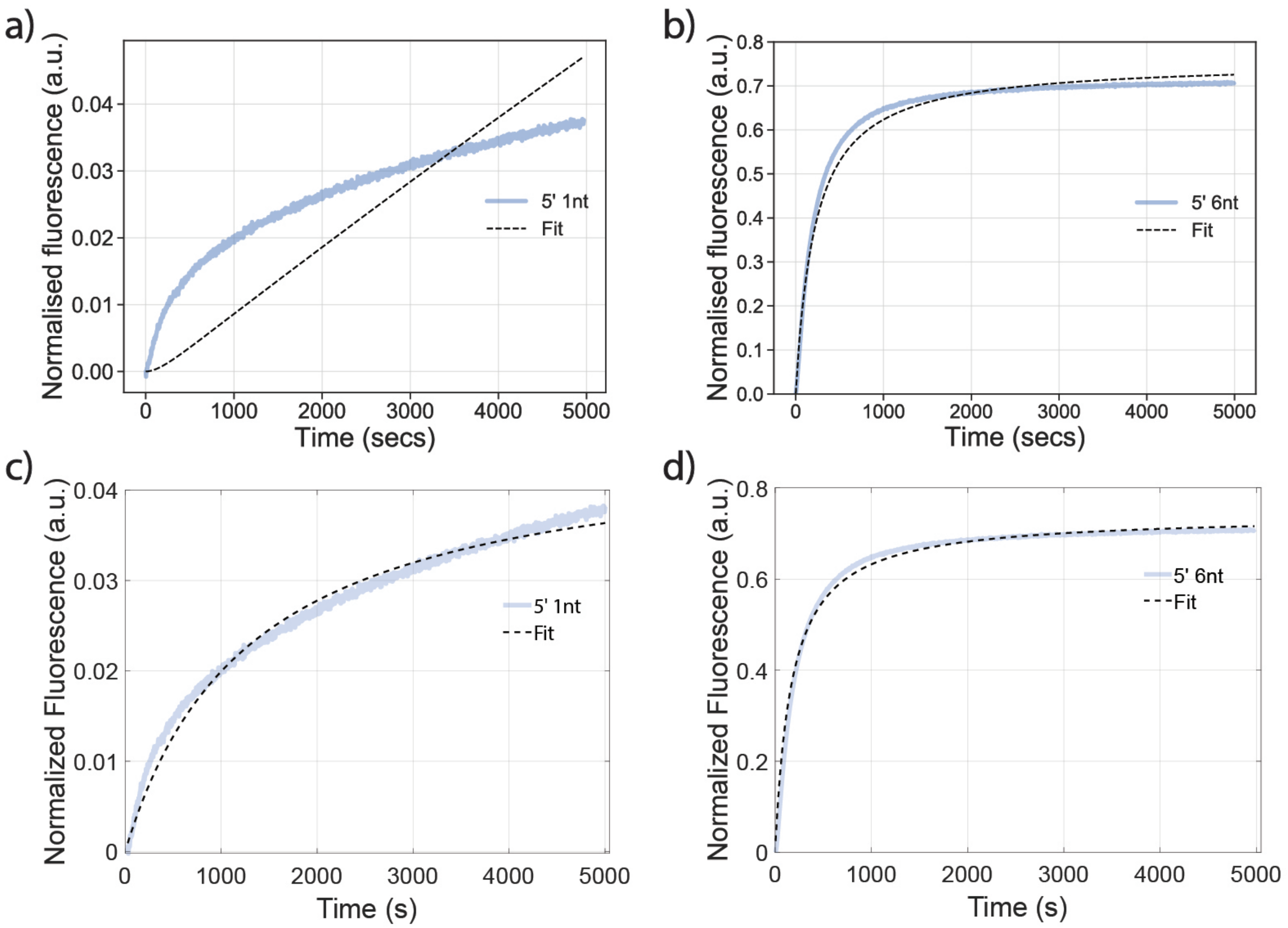


FIG. S7. The fitting result of the experimental data with two methods. All fitting results showing here based on RNA systems with toehold placed 5' and for comparison two reactions are showing: the fastest and slowest one. a). Method I for 1nt toehold; b). Method I for 6nt toehold; c). Method II for 1nt toehold; d). Method II for 6nt toehold.

| Reactions | $\log_{10} k_{\text{rep}}$ | $\alpha$ |
|---|---|---|
| Single 5' | 5.25 (0.08) | 0.86 (0.01) |
| Single 3' | 4.15 (0.03) | 0.91 (0.03) |
| Single DNA | 5.63 (0.03) | 0.85 (0.01) |

TABLE S5. The fitting result of incumbent strand displacing reporter gate with method I. All the rate constants and the associated alpha values are shown as the average of the fitting results of the three replicas and the calculated standard deviation is shown in the bracket.

| Experiment | | | $\log_{10} k_{\text{eff}}$ |
|---|---|---|---|
| RNA→RNA | with 5′ toehold | Toehold 1 nt | 4.10 (0.04) |
| | | Toehold 2 nt | 3.68 (0.03) |
| | | Toehold 3 nt | 4.08 (0.02) |
| | | Toehold 4 nt | 4.61 (0.09) |
| | | Toehold 6 nt | 4.99 (0.06) |
| | | Mismatch P3 | 5.05 (0.03) |
| | | Mismatch P10 | 4.25 (0.01) |
| | | Mismatch P20 | 3.75 (0.01) |
| | | Mismatch P21 | 4.60 (0.01) |
| | | Mismatch P23 | 4.83 (0.02) |
| | | Mismatch P25 | 4.92 (0.03) |
| | with 3′ toehold | Toehold 1 nt | 3.84 (0.15) |
| | | Toehold 3 nt | 3.09 (0.08) |
| | | Toehold 6 nt | 3.94 (0.04) |
| RNA→DNA | with 5′ toehold | Toehold 1 nt | 4.30 (0.16) |
| | | Toehold 3 nt | 3.94 (0.09) |
| | | Toehold 6 nt | 5.17 (0.03) |
| DNA→DNA | with 5′ toehold | Toehold 1 nt | 3.89 (0.07) |
| | | Toehold 3 nt | 3.66 (0.03) |
| | | Toehold 6 nt | 5.23 (0.04) |
| DNA→RNA | with 5′ toehold | Toehold 6 nt | 3.95 (0.04) |

TABLE S6. Estimated rate constants with Method II to the experimental data. All the rate constants are shown as the average of the fitting results of the three replicas and the calculated standard deviation is shown in the bracket.

### SIV. DISCUSSION ON STRAND DISPLACEMENT FREE-ENERGY LANDSCAPE

Based on the thermodynamic parameters predicted using Nearest-Neighbor model by SantaLucia et al[4–6], we develop a model of free-energy landscape of RNAtoRNA, RNAtoDNA, DNAtoRNA, DNAtoDNA TMSD strand displacement. The models of free-energy landcapes are shown in Figure S8. It should be noted that it is not an accurate evaluation of the free energy of the whole process but only a rough estimate, based on the nearest-neighbor parameters.

The free-energy profiles, shown in Fig. S8, start with free-energy decrease until the invader is fully bound to the toehold, as each new base-pair formed decreases the free-energy and increases the stability of the system. The trend of the duplex thermodynamic stability in the nearest-neighbor models is RNA:RNA > RNA:DNA > DNA:DNA. We also include a $2k_{\text{B}}T$ barrier to initiation of the branch migration, as established in Ref. [7]. When RNA displaces RNA (RNAtoRNA) or DNA displaces DNA (DNAtoDNA), the base pair type remains the same (RNA with RNA or DNA with DNA), so we approximate the free energy landscape as flat. For the hybrid systems, however, the existing base pairs between the substrate and the incumbent strands get either replaced by stronger base pair (RNA invades DNA duplex) or weaker base pair (DNA invades RNA duplex), which results in decreasing or increasing free-energy landscape as the branch migration proceeds.

We note, however, that the free-energy landscapes shown in Fig. S8, are very approximative and do not take into

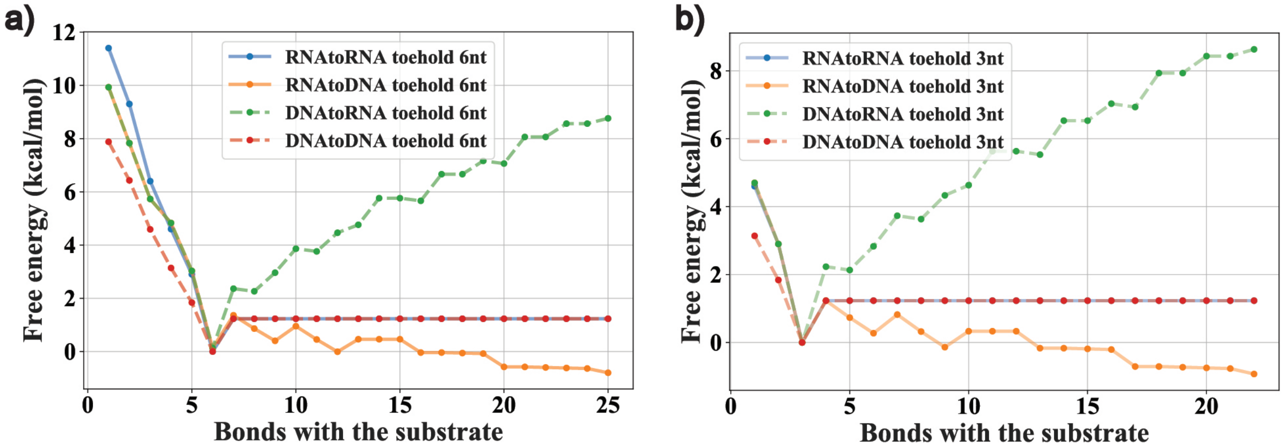


FIG. S8. The thermodynamic profiles for the three strand displacement systems studied with a) The toehold length being 6nt and b) 3nt. For comparison, the point corresponding to the three-way transition state of all three curves is normalized to 0 kcal/mol. Specifically, on the left side of the turning point (corresponding to the transitions state where branch migration is about to start), the green and blue curve overlaps and the red and orange curve overlaps; on the right side, the red and blue curve overlaps.

account e.g. the effects of coaxial stacking interactions. A more detailed free-energy landscapes can be obtained from simulations , e.g. using oxDNA[8] and oxRNA[9]. However, a coarse-grained model of comparable efficiency has not been developed for DNA:RNA hybrids yet. We hence provide our estimated landscape here and note that further research and modeling is needed to study both thermodynamics and kinetics of DNA:RNA hybrids.

---


[1] J. N. Zadeh, C. D. Steenberg, J. S. Bois, B. R. Wolfe, M. B. Pierce, A. R. Khan, R. M. Dirks, and N. A. Pierce. NUPACK: analysis and design of nucleic acid systems. *Journal of computational chemistry*, 32(1):170–173, 2011.

[2] D. Y. Zhang and E. Winfree. Control of DNA strand displacement kinetics using toehold exchange. *Journal of the American Chemical Society*, 131(47):17303–17314, 2009.

[3] R. R. Machinek, T. E. Ouldridge, N. E. Haley, J. Bath, and A. J. Turberfield. Programmable energy landscapes for kinetic control of DNA strand displacement. *Nature communications*, 5:5324, 2014.

[4] J. SantaLucia, H. T. Allawi, and P. A. Seneviratne. Improved nearest-neighbor parameters for predicting DNA duplex stability. *Biochemistry*, 35(11):3555–3562, 1996.

[5] N. Sugimoto, S.-i. Nakano, M. Katoh, A. Matsumura, H. Nakamuta, T. Ohmichi, M. Yoneyama, and M. Sasaki. Thermodynamic parameters to predict stability of RNA/DNA hybrid duplexes. *Biochemistry*, 34(35):11211–11216, 1995.

[6] M. J. Serra and D. H. Turner. [11] Predicting thermodynamic properties of RNA. In *Methods in enzymology*, volume 259, pages 242–261. Elsevier, 1995.

[7] N. Srinivas, T. E. Ouldridge, P. Šulc, J. M. Schaeffer, B. Yurke, A. A. Louis, J. P. Doye, and E. Winfree. On the biophysics and kinetics of toehold-mediated DNA strand displacement. *Nucleic acids research*, 41(22):10641–10658, 2013.

[8] T. E. Ouldridge, A. A. Louis, and J. P. Doye. Structural, mechanical, and thermodynamic properties of a coarse-grained DNA model. *The Journal of chemical physics*, 134(8):02B627, 2011.

[9] P. Šulc, F. Romano, T. E. Ouldridge, J. P. Doye, and A. A. Louis. A nucleotide-level coarse-grained model of RNA. *The Journal of chemical physics*, 140(23):06B614_1, 2014.
